# COMSE: Analysis of Single-Cell RNA-seq Data Using Community Detection Based Feature Selection

**DOI:** 10.1101/2023.06.03.543526

**Authors:** Qinhuan Luo, Yaozhu Chen, Xun Lan

## Abstract

Single-cell RNA sequencing enables studying cells individually, yet high gene dimensions and low cell numbers challenge the analysis. And only a subset of the genes detected are involved in the biological processes underlying cell-type specific functions. We present COMSE, an unsupervised feature selection framework using community detection to capture informative genes from scRNA-seq data. COMSE identified cell substates with high resolution, as demonstrated by its capacity in distinguishing cells at different stages of the cell cycle. Evaluations based on real and simulated scRNA-seq datasets showed COMSE outperformed methods even at high dropout rates in cell clustering. We also demonstrate that by identifying communities of genes associated with batch effects, COMSE differentiates biological differences from batch effects, thereby enabling integrated analysis of scRNA-seq datasets generated with different platforms.

## Introduction

High-throughput single-cell RNA sequencing (scRNA-seq) have enabled the elucidation of transcriptomic profiles on a cell-by-cell basis(1, 2). Statistical and machine learning analyses of scRNA-seq data have provided unprecedented opportunities to elucidate the biological processes underlying phenomena such as cell fate determination(3) and the development and progression of complex diseases (4, 5). However, analyzing scRNA-seq data presents three primary challenges:(a) A typical scRNA-seq experiment detects ∼20,000 genes across just a few thousand cells. Datasets with large numbers of features but relatively small sample sizes are problematic for traditional statistical and machine learning methods. (b) Current scRNA-seq protocols capture only 10-40% of mRNAs within a cell(6–9). As a consequence, lowly expressed genes may be recorded as zeros in the count matrix, and the high dropout rate introduces considerable noise and sparsity into scRNA-seq data. (c) Most functional pathways and biological processes involve only subsets of genes. An important computational challenge is selecting the most relevant features(10). In most cases, true labels for cell types are unavailable for scRNA-seq data; thus, unsupervised feature selection is necessary for both dimensionality reduction and data denoising. Even for high-cell number droplet-based dataset(11), gene selection remains challenging due to extreme sparsity and noise. While sample sizes increase, dropout rates also rise, and most genes are detected in only a tiny fraction of cells (12). Sensitive, scalable methods are needed to distinguish signals of interest. Several computational approaches are commonly applied for gene selection in scRNA-seq data analysis, including Seurat (13), M3Drop (10) scran(14), BASiCS (15), and scLVM (16). These methods share two main components: data normalization and analyzing variation. Normalization is often achieved through variance-stabilizing transformation, as in DESeq2 (16), or converting raw counts to relative expression. Different approaches were adopted to highly variable genes by these methods. For example, scLVM’s LogVar algorithm and scran calculate variance from a logarithmically normalized expression matrix. BASiCS uses hierarchical Bayesian model, while Brennecke uses the squared coefficient of variation (CV2) to estimate dispersion in expression. Each method fits mean-variance relationships to their model and selects highly variable genes. Identification of highly variable genes (HVGs) improves signal to noise and computational efficiency of downstream tasks like clustering by reducing genes considered. However, HVGs often correlate or are redundant, especially in heterogenous populations. Redundancy can cause overfitting, variance inflation, low efficiency and poor performance. Moreover, not all biologically relevant genes are highly variable, so selection based solely on mean-variance relationships causes information loss.

Feature selection aims to identify the most informative and relevant features for building models in tasks such as classification, trajectory analysis(17), or other downstream analyses. It also facilitates model interpretability and generalizability. We focus on unsupervised feature selection (UFS) methods, as true cell type labels are often lacking in scRNA-seq data. We proposed a novel unsupervised feature selection (UFS) method named as COMSE. This method first partitions all genes into different communities in latent space inferred by Principle Component Analysis (PCA) using the Louvain algorithm (18). Within each community, we apply a denoising procedure to remove noise introduced during sequencing or other procedures. It then selects highly informative genes from each community based on the Laplacian score (19) (Fig1). Louvain algorithm is a hierarchical clustering method that optimizes modularity to detect community structure. Laplacian score is a feature selection approach based on spectral graph analysis that ranks features by local neighborhood connectivity and global distinctiveness using similarity matrix.

**Figure 1.**
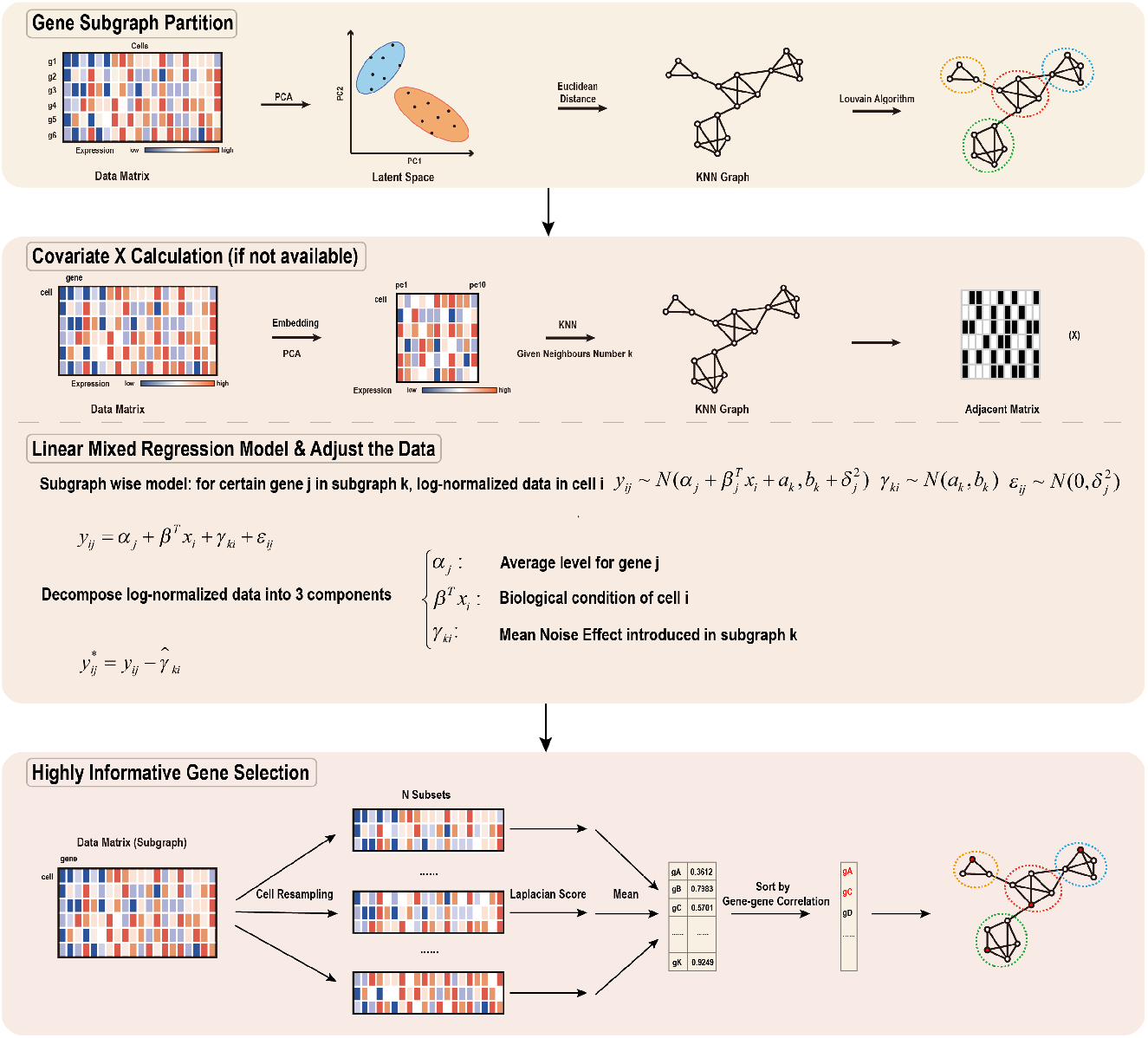
An overview of the COMSE method for selecting informative genes from single-cell RNA-seq data. (A) The log-normalized gene expression profile of each cell is projected into a low-dimensional latent space using principal component analysis (PCA). The low-dimensional representation is then used to construct a gene similarity graph with the K-nearest neighbors (KNN) algorithm. Gene graph is partitioned into several subgraphs by Louvain algorithms for community detection. (B) When sample covariates lacking, covariate matrix for each cell were estimated through KNN in low-dimension PCA space with given neighbor number. Then linear mixed regression model was applied to estimate and remove noise from data within each gene subgraph. (C) An unsupervised feature selection technique based on the Laplacian score is applied to choose highly informative genes in each subgraph.

To assess the performance of our method COMSE, we applied it to actual and simulated scRNA-seq data. We found that COMSE was more sensitive to detect subtle differences between homogenous cells without any extra information, which facilitates a more sophisticated understanding of the functionality inherent to cellular subpopulations (Fig2). Moreover, COMSE gave more precise and concise cell clustering than other commonly used tools (Fig3, Fig S2,3). We also demonstrated that the communities identified by our method provide insight into gene function at both broad and gene-specific levels (Fig4A, B). The community structure is also able to separate subgroups associated with batch effects, allowing us to remove such effects (Fig4C, D). Therefore, we show that COMSE could aid in interpreting variability arising from either biological or technical sources. We also applied the denoising step in COMSE to bulk RNA-seq data, the results showed that after denoising, the data yielded a more robust identification of differentially expressed genes and better reflected the biological differences between groups (Fig5).

**Figure 3.**
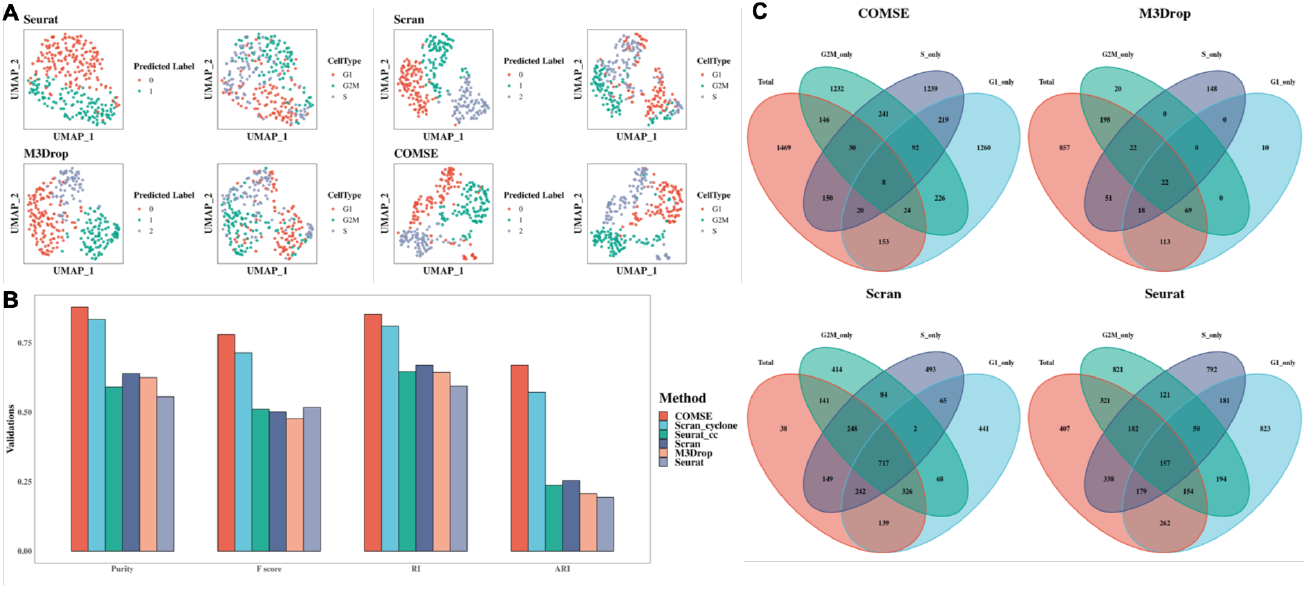
COMSE identified cell-cycle states in mouse embryonic stem cells (ESCs). (A) The UMAP plots of the real scRNA-seq data for cell-cycle states of mouse ESCs grouping by predicted label using genes selected by different feature selection methods and true label. (B) Four external clustering validations (Purity, F score, RI, and ARI) were used to indicate the performance and robustness of the four methods with the mouse ESCs dataset. (C) The Venn plots of genes selected by each feature selection method using G1 phase mouse ESCs only, G2M phase mouse ESCs only, S phase mouse ESCs only, and all mouse ESCs.

**Figure 3.**
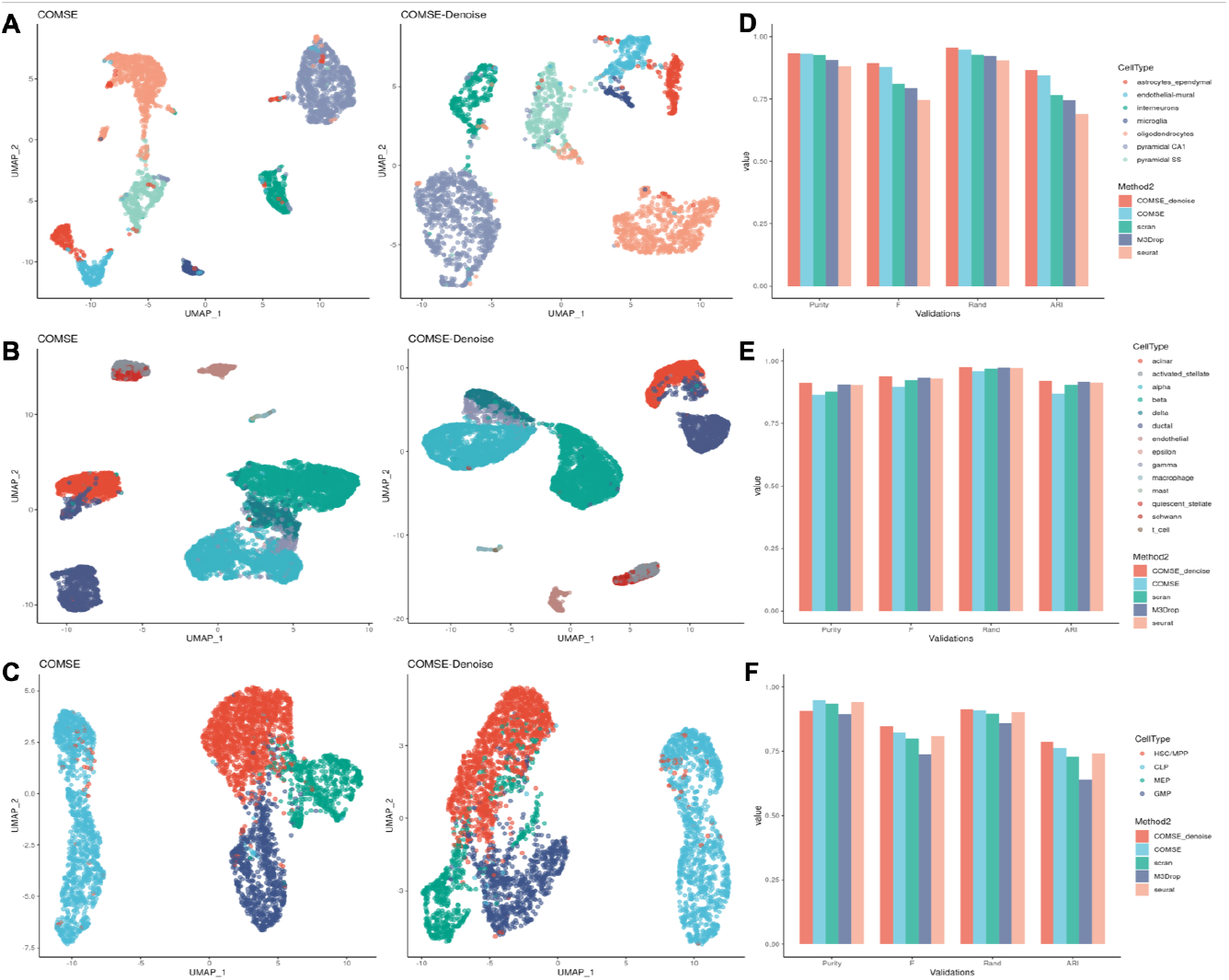
COMSE yields more accurate cell classification results with four scRNA-seq datasets. (A, B, C) UMAP plots of the mouse brain dataset, Baron pancreas dataset and Bunis HPSC dataset using top 2000 genes obtained from COMSE with denoising, COMSE only and other three widely used HVG selection methods. (D, E, F) The cell clustering results using top 2,000 genes selected by COMSE with denoising, COMSE only and other three widely used HVG selection methods of three heterogeneous public scRNA-seq datasets with true labels, including a mouse brain cortex (Zeisel) dataset, the human pancreas (Baron) dataset and human hematopoietic stem and progenitor cell (Bunis HPSC) dataset. Four external clustering validations (F score, CSI, PSI, ARI, and NMI) were used to indicate the performance and robustness of the five methods on three public scRNA-seq datasets.

**Fig4.**
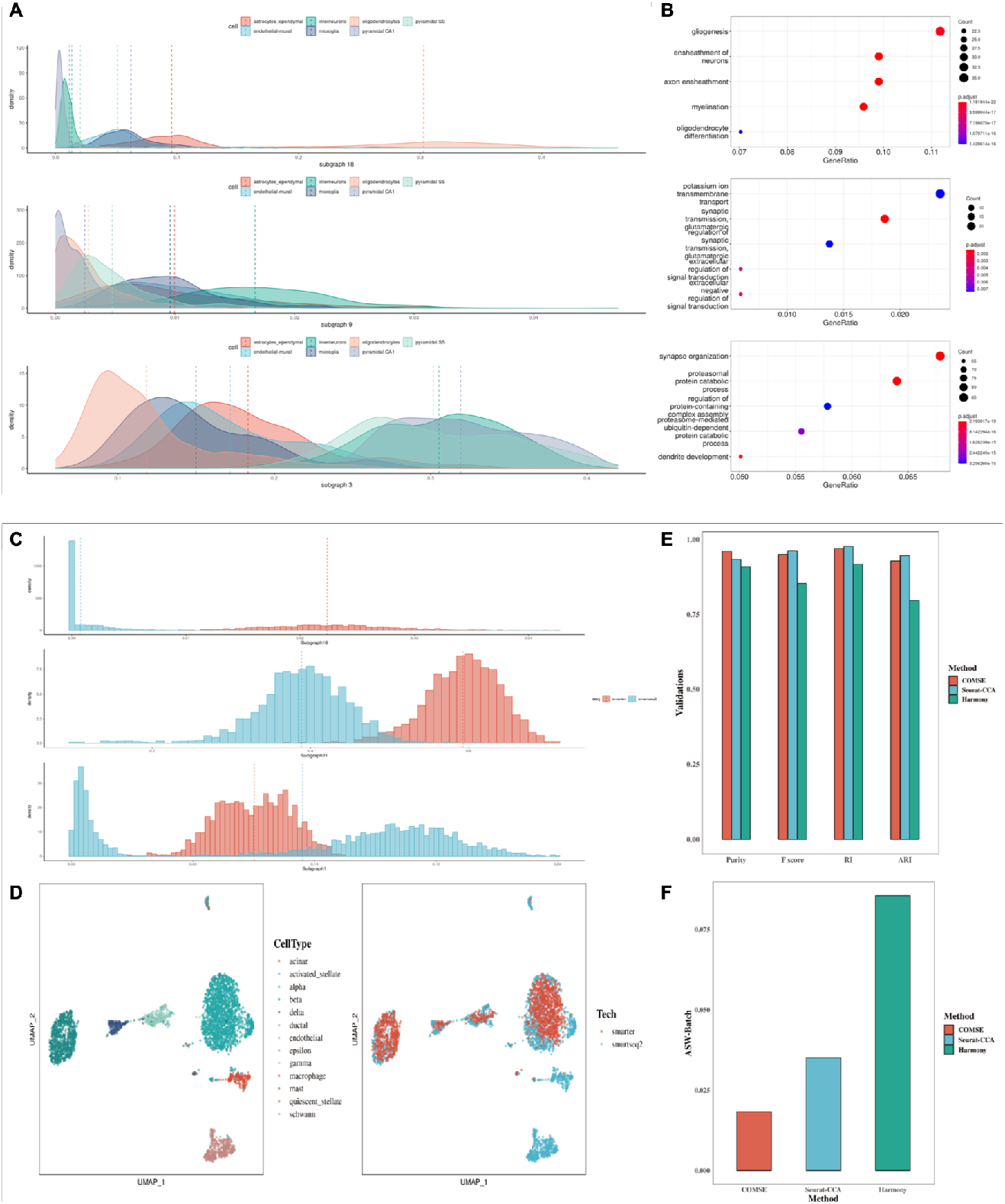
Biological interpretation of subgraph generated by COMSE. (A) Density plot of the results obtained with mouse brain dataset and of the activities of each subgraph in each cell generated by COMSE. The different color represents heterogeneous cell populations. (B) Dot-plot of the results obtained by introducing gene ontology (GO) pathways analysis into subgraphs 3, 9, 18, only the top 5 categories were shown. COMSE removed the batch effect introduced by different scRNA-seq protocols with high efficiency using two human pancreatic islet datasets. (C) Histogram of the results obtained with the human pancreatic islet dataset and of the activities of each subgraph in each cell generated by COMSE. The red color represents the cell using the scRNA-seq protocol SMARTER while the blue color represents the cell using SMART-seq2. (D) After using the AUC scores of three subgraphs to regress out the highly informative genes with high variability selected by COMSE, UMAP plots of the human pancreatic islet datasets were obtained with the top 2000 genes selected by COMSE. (E) Four external clustering validations (Purity, F score, RI, ARI) were used to indicate the performance of COMSE on cell clustering. (F) Average Silhouette Width (ASW) assessments of different batch effect removal methods on the human pancreatic islet dataset, Smaller values of which indicate better batch effect removal and less heterogeneity introduced by different protocols.

**Fig5.**
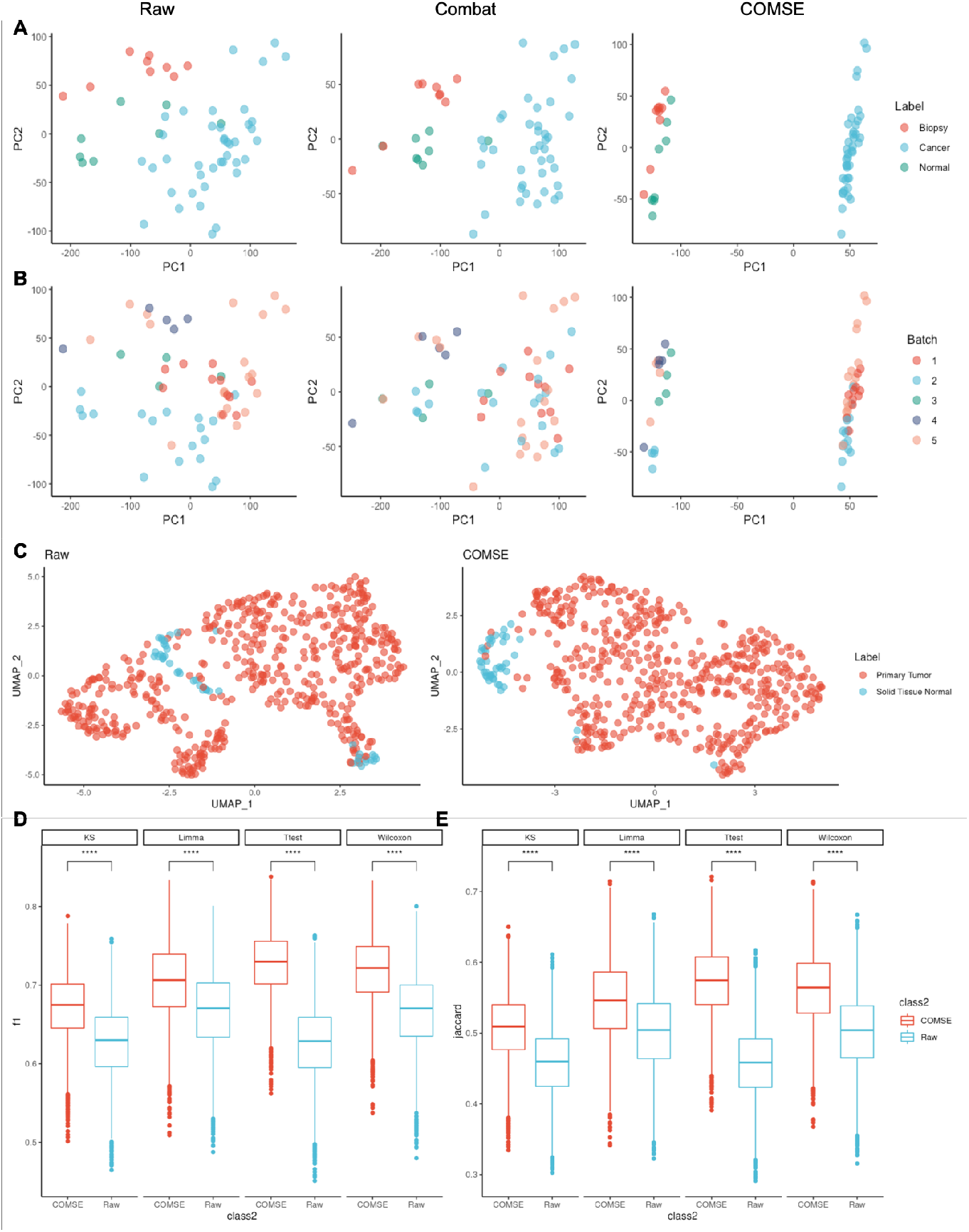
(A, B) The PCA plots of the bladder bulk RNA-seq original data, ComBat-modified data and COMSE-modified data, denoted by sample type and batch. (C) The UMAP plots of the native TCGA HNSC data and COMSE-processed data, labelled by sample type. (D) The Boxplots of paired F1 values computed from differentially expressed genes (DEGs) extracted via each approach including t-test, Wilcoxon-test, limma and KS-test in each group. P-values were derived using the t-test. (E) Boxplots of paired Jaccard similarity coefficients between DEGs elicited via each approach including t-test, Wilcoxon-test, limma and KS-test in raw down-sampled data collection and denoised down-sampled data collection separately. P-values were produced using the t-test.

## Materials and methods

### The workflow of COMSE

We proposed an approach that consists of three main steps (Figure 1). First, we construct a K-nearest neighbor graph (KNN graph) based on the Euclidean distance between genes in the low-dimensional space obtained by principal component analysis (PCA). Then, we partition the KNN graph into multiple subgraphs using the Louvain algorithm for community detection.

Secondly, we perform a denoising procedure for heterogeneous gene expression datasets. For scRNA-seq data, where covariate information is unavailable for each cell, we first infer a covariate matrix based on a KNN graph within the low-dimensional space from PCA. We then implement a linear mixed regression model to remove noise introduced by different methods in each subgraph. For bulk RNA-seq data, we can extract sample group labels (e.g. “Cancer” vs. “Normal”) to use as covariates. Thus, inferring a covariate matrix is unnecessary. In addition, when examining scRNA-seq datasets for cells with similar properties or in high quality, this denoising step is optional.

Finally, we select the informative genes in each subgraph through the Laplacian score. For each gene in a subgraph, we calculate the Laplacian score with multi-subsample randomization and choose genes with the smallest scores, assuming that data from the same class are often close to each other. We then rank the genes based on gene-gene correlation to remove redundancy.

### Data preprocessing and normalization

scRNA-seq data is always organized as a data matrix with rows as genes and columns as cells, which is the input of COMSE. We only kept genes in count matrix expressed more than three cells. Each gene expression values for each cell were divided by the total values of each gene and multiplied by 10,000. These normalized values were then natural-log transformed using log1p function before further downstream analyses.

### Gene graph construction

We first introduced principal components analysis (PCA) into log-normalized data matrix, using the prcomp function in R, after scaling and centering the data. To overcome the extensive technical noise in any single cell for scRNA-seq data, we selected the number of PCs to keep through the ‘elbow’ according to Elbow-plot (a ranking of PCs based on a percentage of variance explained by each PC). In all datasets used in our studies, we chose top 3 PCs for downstream analyses. After we embedded the original log-normalized data matrix into low-dimensional latent space inferred by PCA, a K-Nearest-Neighbors (KNN) gene graph was constructed based on the Euclidean distance in latent space with given K (configurable, but typically around 30, we used 30 here, a common value for graph construction(20, 21)) neighbors. Then we binarized the KNN graph by assigning non-zero edge to 1 and this binarized KNN graph was used as the adjacent matrix between genes for the following analysis. And this binarized adjacent matrix constructed gene graph.

### Gene subgraph partition

Louvain algorithm is a widely used community algorithm based on the modularity and a hierarchical approach with the goal of maximizing the modularity. This algorithm can be used for partitioning undirected graph to find relevant nodes in graph without overlapping. We performed Louvain algorithm for gene graph and obtained dozens of subgraphs partitioned from binarized adjacent matrix using multilevel.community function in R. We supposed that we could partition the whole gene graph into several functional modules which could improve our interpretation of variability introduced by heterogenous cell populations or different batches. Through this step, we obtain a series of subgraphs with functional potential. More importantly, for each subgraph, the quantity difference between cells and genes becomes smaller, or the number of cells even exceeds the number of genes. In this way, we have largely addressed the imbalance between features and sample sizes.

### Covariate X calculation

We first re-normalized the scRNA-seq data matrix by scaling each gene’s expression values for each cell by the total expression values of that cell and multiplying by 10,000. The re-normalized values were then log-transformed using the log1p function. We performed PCA on the normalized and log-transformed data, retaining components that capture major variability. We used the PCA space to build a KNN graph connecting cells with similar expression patterns, using 30 nearest neighbors by default. We binarized this KNN graph by setting all non-zero edge weights to 1. The binarized KNN graph served as an adjacency matrix between all cells and was then utilized as the covariate matrix X in our subsequent denoising procedure. For bulk RNA-seq data, sample group labels could be used as covariate X directly.

### Linear mixed regression model for denoising

We assume that noise affecting genes within the same subgraph and cell, from various sources, follows a normal distribution with the same parameters. Then, we use a linear mixed regression model to decompose gene expression in each cell and subgraph into baseline average expression *α*_*j*_, biological condition of cell i, and mean noise components *γ*_*ki*_. By estimating the mean noise (*γ*_*ik*_) for each cell in a subgraph, we obtain a proxy for the total nuisance technical variation, allowing us to denoise the data.

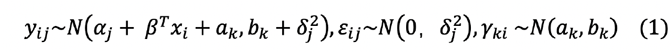

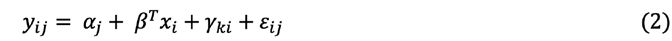

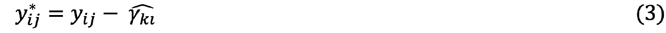

### Highly informative gene selection

We chose highly informative genes in each subgraph independently. For each subgraph, we first randomly selected n cells (configurable, we used 100 to 300 here based on the number of cells in the dataset) with replacement N (default 100) times. We then calculated the Laplacian scores for each gene in each subset. We calculated the mean Laplacian score for each gene across all subsets as the final Laplacian score for each gene. However, we found that the correlations among the top genes were usually large, meaning that directly selecting the top genes would lead to information redundancy. To address this issue, we ranked the genes using the following steps based on gene-gene correlation (Algorithm 1).

#### Algorithm 1 Gene Sorting (subgraph — wise)

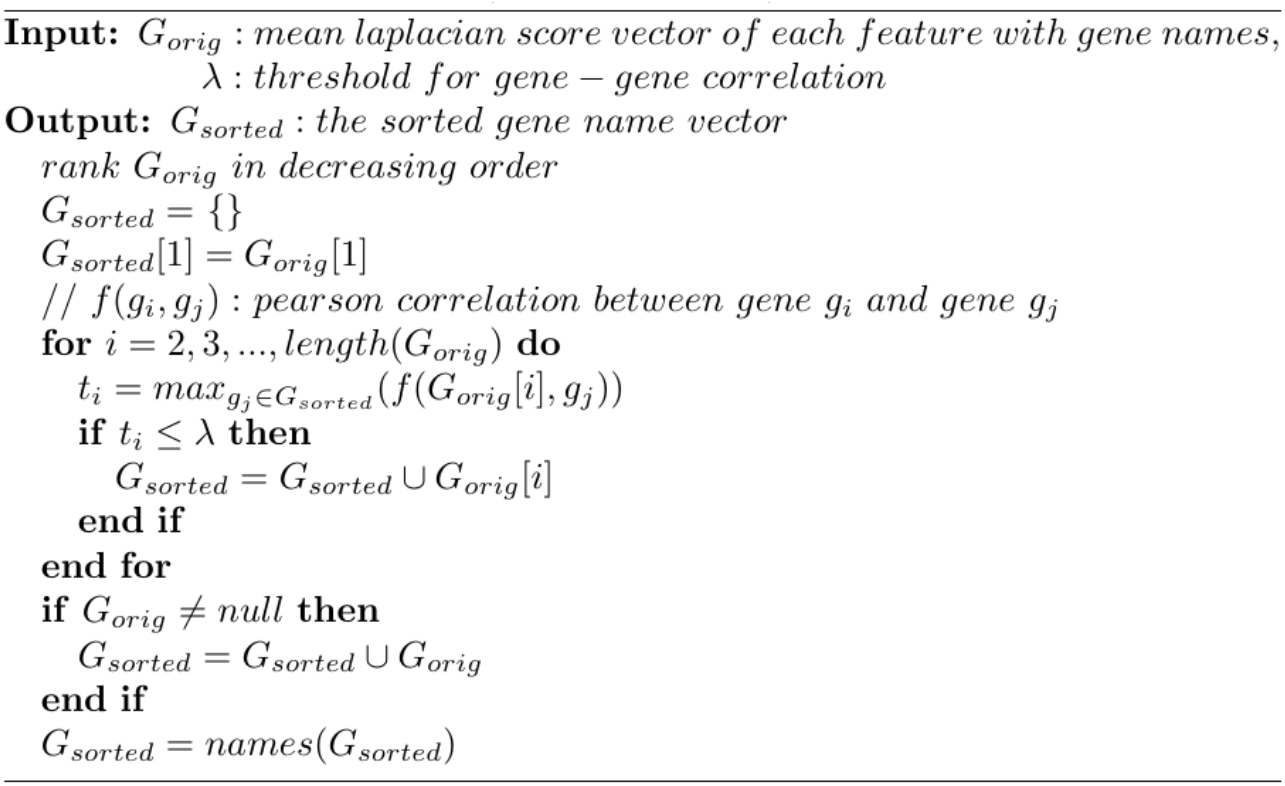

Specifically, when analyzing data with lower cellular heterogeneity, such as PBMC data, we set to between 0.3 and 0.5. When analyzing data with substantial biological variability introduced by heterogeneous cell populations, we set to 0.5. Once we obtained for each subgraph, we selected highly informative genes in equal proportions based on the number of genes in each subgraph.

### Cell classification

After obtaining the highly informative genes selected from COMSE or highly variable genes from other widely used methods, we used these genes as input for PCA reduction using the RunPCA function in Seurat. We then used the first 10 principal components as input for cell classification, UMAP reduction, and visualization following the standard workflow of Seurat for analyzing single-cell RNA-seq data. To determine the optimal resolution for clustering, we tested 10 different resolutions (from 0.1 to 1) for each feature selection method and evaluated the results using four external validation metrics. The resolution with the best overall scores across these metrics was selected for further analysis of the data processed by that method.

### AUCell activity score

We can quantify the activity of subgraphs in each cell using AUCell (22). AUCell takes gene sets of interest as input and converts the expression matrix of genes in cells into an AUC matrix of subgraph activity for each gene. We used the AUC activity score in batch effect-associated subgraphs as covariates to remove batch effects.

### Batch effect removal

We first calculated the AUC activity score of subgraphs 1, 16, and 21 obtained by the Louvain algorithm. We then identified the top 2,000 highly informative genes (HIGs) selected by COMSE and the top 2,000 highly variable genes (HVGs) selected by Seurat. We considered the intersecting genes between HIGs and HVGs to be highly informative genes with high variability. We hypothesized that these highly informative genes with high variability were more likely to reflect systematic variability introduced by different batch effects. Therefore, we used the ScaleData function in Seurat with a ‘negbinom’ model to regress out these highly informative, highly variable genes using the AUC scores of subgraphs 1, 16, and 21. We then performed standard single-cell RNA-seq analysis and evaluation using Seurat, including cell clustering, UMAP reduction, and visualization (Figure 4D, S4,6).

### Cluster validation

External indexes use the true label for comparison, among the hundreds of clustering validation methods, we selected some widely used indexes that consider external validation for reference, including Purity, F score, Rand Index (RI), and Adjusted Rand Index (ARI). Purity measures the ability of a clustering method to recover known classes. F score is the harmonic mean of precision and recall. RI measures the similarity between two known clusters. ARI is adjusted for the chance grouping of elements.

### Average silhouette width (ASW)

We performed the ASW (23) on the batch labels to assess batch correction. The silhouette score of each cell is calculated by subtracting its average distance to other members in the same cluster from its average distance to all members of the neighboring clusters, and then dividing by the maximum of the two values. The range of result score is between -1 and +1, where a high score means that the cell fits well in the current cluster, while a low score denotes a poor fit. The average score of all cell can be used for evaluating overall batch mixing through batch effect labels, for batch effect, the smaller the better. In our study, we used the UMAP embedding of the datasets as input to calculate the distance among cells to obtain the ASW scores.

### Bulk RNA-seq data analysis

During bulk RNA-seq data assessment, sample group labels comprise the Covariate X matrix. Consequently, subsequent to subdividing gene subgraphs, we estimated the mean noise effect of each subgraph’s according to Equation (2), thereby procuring denoised data.

We performed PCA on the original and denoised bladder bulk RNA-seq data(24) and visualized the first two principal components. For TCGA HNSC bulk RNA-seq data (25), we first selected the top 2000 highly variable genes following the standard Seurat protocol. We then chose the first 10 principal components for UMAP dimensionality reduction and visualization of the original and denoised data.

To compare the reproducibility of DEGs detected from subsample data before and after noise removal, we repeatedly down-sampled 100 samples from the full HNSC data for 1000 times. For each down-sampled data, after removing noise, we used four methods (limma(26), t-test, Wilcoxon test, and KS test) to screen DEGs in both the raw and denoised subsample data. We directly used limma to find DEGs between normal and tumor groups. For t-test, Wilcoxon test, and KS test, we first calculated p-values based on each gene’s expression in the tumor versus normal groups. Then we adjusted the p-values using the Benjamini-Hochberg (BH) method for multiple testing correction. We considered any gene with an adjusted p-value less than 0.05 as a DEG for further analysis. To evaluate DEG reproducibility before and after denoising, we calculated the paired Jaccard similarity and F score between DEGs identified using each method in the 1000 original and denoised down-sampled datasets separately.

### Jaccard similarity

The Jaccard similarity index quantifies the resemblance between two sample collections, defined as the cardinality of their intersection divided by the cardinality of their union. Its values span 0 to 1, where 0 signifies the two sample groups completely differ and 1 denotes they perfectly agree.

## Results

### COMSE identifies sub-states in one cell population

Approaches for highly variable gene selection typically filter for features (genes) exhibiting greater variance, specifically larger biological variance. These techniques frequently enhance downstream analyses like cell clustering and pseudo-time inference. However, differences between subtypes of the same cell type are commonly minute. Variance within a cell type may subsume variance between subtypes, confounding the separation of in-group variance and between-group variance. Consequently, unambiguously discriminating distinct subtypes of the same cell type may prove challenging.

To further validate our hypothesis, we generated a dataset containing three groups and three types of features. The first type of feature was sampled from three normal distributions for different groups, with small mean differences between the three distributions and small variances. The second type of feature was also sampled from three normal distributions for different groups, with similarly small mean differences but larger means and smaller variances. The third type of feature was sampled from a single normal distribution for all three groups, with a mean similar to the second feature type and larger variance than other types of features. Only the first two types of features represent between-group differential genes for each group, while the last feature type corresponds to the in-group differential genes we mentioned above. We found that inter-group differential genes can effectively separate the three sample groups. Furthermore, we used feature selection methods to screen for features for downstream clustering analysis. However, differentially expressed genes selected based on commonly used HVG detection methods had difficulty distinguishing the three sample groups, while the features selected by our method separated the three groups well, further validating our hypothesis (FigS1).

Single-cell RNA sequencing (scRNA-seq) is commonly used to identify different cell types from heterogeneous tissues or complex cell populations. However, even seemingly one cell population consist of cells in distinct cellular states due to differences in developmental stage, cell cycle phase, or microenvironment. Several studies have found that the cell cycle contributes to phenotypic and functional cell heterogeneity even in homogeneous cell populations(27, 28). Due to the high prevalence of dropout events and technical noise in scRNA-seq data, it is difficult to distinguish cells in different phases of the cell cycle in a homogeneous cell population (27).

Therefore, we analyzed a scRNA-seq dataset of mouse embryonic stem cells (ESCs) sorted into the G1, S and G2M phases of the cell cycle. Cells in different phases were not clearly separated using the top 2,000 highly variable genes selected by widely used HVGs selection methods with the standard Seurat workflow (Fig2A). We hypothesized that the selected HVGs are in-group differential genes rather than between-group differential genes. To verify this, we applied highly variable genes (HVGs) selection methods separately to cells in G1, S and G2M phases. We observed extensive overlap among the highly variable genes selected from cells in different cell cycle phases (Fig2C). This suggests that HVGs selection methods may have difficulty differentiating the relatively subtle differences in gene expression that distinguish distinct cellular states within a identical population. While highly variable gene selection can be useful for identifying major sources of variation, more nuanced methods may be needed to resolve minor differences within a seemingly homogeneous cell population. Using the top 2,000 genes selected with our proposed method, COMSE, we generated a clear cluster structure reflecting the different cell cycle states with the highest performance across four external clustering validation methods.

Currently, there are also methods for distinguishing different cell cycle phases, such as the cell cycle score in Seurat and Cyclone in scran. These methods generally classify cells based on known cell cycle-related genes, i.e., by pre-specifying between-group differential genes. We also included both these methods in our comparison. Overall, we found that our method performed better in distinguishing cells in different cell cycle phases (Fig2B). Therefore, our method has higher sensitivity for distinguishing different subtypes of the homogeneous cell type.

### Accurate and robust single-cell classification using COMSE

To evaluate the performance of our proposed method on cell clustering analysis, we collected single-cell RNA-seq datasets with true labels, including the data from the cerebral cortex of mouse(29) (GSE60361, 3005 single cells from 33 males and 34 females), the dataset from the pancreas of human(30)(GSE84133,8569 single cells containing 14 common cell types in pancreas), the dataset from the human fetal, newborn, and adult haematopoietic stem and progenitor cells (31)(GSE158493, 5183 single cell samples containing 5 cell types), the dataset Zhengmix4uneq (32) containing four pre-sorted cell types (1,000 B-cells, 500 naive cytotoxic T-cells, 2,000 CD14 monocytes and 3000 regulatory T-cells) from Homo sapiens, the dataset mixology10×5cl containing five human lung adenocarcinoma cell lines (GSE118767, 3918 cells from H2228, H1975, A549, H838, and HCC827), the dataset Koh (33) containing purified H7 human embryonic stem cells (GSE85066, 531 cells from 9 FACS purified differentiation stages). As previously articulated, the denoising procedure predominantly proves pertinent for interrogating heterogeneous corpora. Accordingly, within our analyses, we enlisted COMSE and COMSE with denoising step for the Bunis HSPC data, Zeisel brain data, and Baron pancreas data. For residual arrays, we eschewed the denoising maneuver. In addition to COMSE, we utilized the widely used M3Drop, Scran and Seurat for cell clustering comparison.

Primarily, we extracted the top 2000 feature genes from COMSE with denoising, COMSE alone, scran, M3Drop, and Seurat. Cell clustering was ensued in compliance with the Seurat conventional procedure. To evaluate the performance of cell clustering, we enlisted four external clustering validation implements (Purity, F score, RI, ARI).

Amid the Zeisel brain dataset and Baron pancreas dataset, all four clustering validations systematically evinced COMSE with denoising accomplished best cell clustering. As for the Bunis HPSC dataset, COMSE with denoising also showed the better performance on F score, RI and ARI and similarly to other common HVG selection methods in purity (Fig2D, E, F). For other simpler data (with fewer cell types or bigger differences between heterogeneous cells), we also found that COMSE gave accurate clustering results (FigS2). Furthermore, to validate the performance of our method with different numbers of features, we compared the clustering results using top 500, 1000, and 2000 features. We found that our method achieved robust performance under different feature numbers, indicating that our method can not only obtain more accurate cell clustering results but is also robust to the number of features (FigS3).

### COMSE enhances the interpretation of variability

As the delineation of gene subgraphs depends upon Euclidean separation in the latent space extrapolated by PCA, we speculated genes inhabiting the equivalent subgraph may represent a functional module. We used subgraphs partitioned from gene graph through the Louvain algorithm using a mouse brain dataset to demonstrate functional relevance. We calculated the AUC activity score of each subgraph in each cell generated by COMSE. Then we selected subgraphs having differential activity scores among heterogeneous cell populations, which were subgraphs 18, 9, and 3 (Fig4A, B). The Gene Ontology (GO) analysis of genes in these three subgraphs demonstrated the consistency between the enriched signal pathways using genes in each subgraph and the function of cell types that have exclusively high AUC scores in the corresponding subgraph. Oligodendrocytes are the myelinating cells of the central nervous system, whose main function is to generate myelin, which is an extended membrane from the cell that wraps tightly around axons (34). And what contains genes enriched in gliogenesis and myelination pathways are highly consistent with the function of oligodendrocytes in the central nervous system which had exclusively high AUC scores in subgraph 18 (Fig4A, B). Besides, interneurons had exclusively high AUC scores in subgraph 9 which contained genes enriched in pathways relative to signal transduction and transmission of neurons. And neurons including pyramidal CA1, pyramidal SS, and interneurons had activity scores in subgraph 3 which contained genes enriched in the pathways involved in the function of neurons such as synapse organization and dendrite development (Fig4A, B). Above all, we verified our speculation to a certain extent, the subgraphs partitioned from the gene graph could be identified as functional modules and could improve our understanding of biological variability among heterogeneous cell populations.

Due to the development of fast and accurate scRNA-seq technologies, the number of cells and studies using scRNA-seq technology grows rapidly (35). Another challenge in single-cell transcriptomics data analysis is the batch effect, which is a systematic bias of transcriptomics profile introduced by one batch to another such as different scRNA-seq protocols (36). The systematic variability generated by different batches could confound biological variations. Therefore, beyond the biological variability, we also wonder whether subgraphs generated by COMSE could be used for removing systematic variability introduced by batch.

We combined two publicly available datasets on the human pancreas generated using SMA(37), and SMART-seq2(38). Before the batch effect removal, the biological variability of heterogeneous cell populations was covered by the systematic variability introduced by different scRNA-seq protocols (FigS4). Since the outcome of COMSE in step 2 was the subgraph partitioned from the whole gene graph through the Louvain algorithm, we were curious whether we could find out some specific subgraph that was batch-relevant. We first found the top 100 differential genes between cells from two different scRNA-seq protocols. These genes enriched in subgraphs 1, 16, and 21 exclusively (FigS5). Then we calculated the AUC activity score of subgraphs 1, 16, and 21 in each cell using the AUCell R package (22). Surprisingly, the AUC activity score of these three subgraphs could separate the cell from two scRNA-seq protocols (Fig4C). Therefore, we supposed these subgraphs could explain the systematic variability introduced by protocols, then we scaled the original data matrix by regressing out all the AUC scores of three subgraphs, and the scaled data was used for downstream analyses. After using the AUC score to regress out highly informative genes with high variability, we could observe that the batch effect introduced by protocols was removed efficiently with no large impact on cell clustering assignments (Fig4D). Besides COMSE, we utilized the widely used Harmony and Seurat CCA for batch effect removal comparison. The cells from two scRNA-seq protocols integrated well in each cell type cluster through Harmony, but the scRNA-seq dataset using SMARTER did not contain any acinar and ductal cells while the integrated data have shown some cells of SMARTER merged acinar and ductal cells from SMART-seq2 (FigS6). To systematically evaluation of the efficiency of batch effect removal and the accuracy of cell clustering, we used Purity, F score,RI, ARI and ASW five widely used metrics. COMSE outperformed other widely used methods with these two validations (Fig4E, F). Above all, COMSE was shown to enable a highly interpretable and efficient batch effect removal.

### Harnessing denoising in COMSE elevates bulk RNA-seq Inquiry

Bulk RNA-seq data analysis can be substantially hindered by considerable noise or batch effects arising from discrepancies in sampling locations, operators, and data origins. When replicate samples are available, batch effects can be eliminated using techniques such as ComBat(39). However, for certain public databases like TCGA, most samples lack replicates, complicating the removal of noise intrinsic to TCGA data. Our denoising approach operates on a sample-wise basis, obviating the need for replicates across all samples. Consequently, our methodology can attenuate noise from both data with and without replicates, demonstrating robust reproducibility.

To substantiate this, we first deployed the denoising procedure on bladder data from bulk RNA-seq. We ascertained the processed data could adeptly distinguish discrete biological conditions while congregating biopsy and normal tissues belonging to the normal group, indicating no supplementary noise was introduced (Fig5A, B).

Crucially, our denoising maneuver functions independently of replicates, circumventing limitations faced by existing batch effect correction techniques reliant on replicate concordance. By operating at the sample level, our approach exhibits marked resilience and wider applicability across diverse data types, especially data lacking replicate samples such as TCGA. Therefore, we analyzed data from TCGA HNSC samples and found the denoised data could better distinguish different biological conditions (Fig5C, D).

More importantly, in routine experimental designs, we may not have large sample sizes like in the TCGA database. Therefore, we hope to obtain relatively stable estimates from a small number of samples. For bulk RNA-seq data analysis, stable estimation often means the identified DEGs show strong consistency. Therefore, we downsampled the TCGA HNSC data by sampling 100 samples from the data each time and selected DEGs between cancer and normal groups using t-test, Wilcoxon-test, KS-test, limma(26), repeating 1000 times. We then compared the similarity of DEGs obtained from each group of samples by calculating the pair Jaccard similarity and F-score of the DEG lists. We found DEGs obtained after denoising step processing showed higher similarity while the number of DEGs obtained from the data did not differ much from the original data (Fig5D, E, FigS7). That is, the denoising step did not significantly affect the power to screen for DEGs in the data.

In summary, our method can not only better reflect differences between biological conditions but also enable more robust analysis results.

## Discussion

Many studies have shown different methodologies for highly variable gene selection in analyzing single-cell transcriptomics data. These widely used methods achieved dimensional reduction as well as denoising. However, it’s difficult for these methods to distinguish the source of variability in some scenarios, especially when variability introduced by biological differences and other batch effects are similar in amplitude. As we analyzed the identical cell population at different cell-cycle stages, it was difficult to distinguish cells from different stages using highly variable genes (Fig2A).

In this paper, we proposed a new approach for feature selection based on community detection, named COMSE. Using this approach, we could acquire highly informative genes from different subgraphs, even genes in some subgraphs were functionally conserved, which were not highly variable. And we could use these highly informative genes to distinguish different cell-cycle stages in the same cell populations (Fig2A, B). Besides, we could speculate that genes in the same subgraph have a similar function, in other words, the Louvain algorithm partition the whole gene graph into several functional modules (Fig4A). Furthermore, these subgraphs could be regarded as the source of systematic variation introduced by different modules, such as different functional modules and other potential batch effects, which meant that we could use these subgraphs for variation decomposition. To verify this speculation, we also collected single-cell RNA-seq data using different protocols, after regressing out the highly variable genes in highly informative genes using AUC scores calculated from three batch-related subgraphs (Batch effect removal in Methods), we could remove the batch effect introduced by different protocols using this highly interpretable and efficient strategy (Fig4C). These comprehensive experiments showed that we were free to remove the variability introduced by unconcerned covariates on the data or figure out the main effect of these covariates on the data which could provide new insights into uncovering the relationship between covariates and different cell populations.

Moreover, when we analyzed the single-cell RNA-sequencing data, we were concerned about the quantity of selected feature genes for downstream analyses. COMSE shows the robustness in performance of cell clustering using the different numbers of highly informative genes. We mainly attribute the robustness and high accuracy in cell clustering of COMSE to the highly informative gene selection strategy applied in each subgraph separately. First, we selected genes from each subgraph in equal proportions. Therefore, using these highly informative genes even with small numbers, we could obtain a more complete depiction of data, which allowed us to get better estimates even with less information. Also, the denoising procedure ameliorates bulk RNA-seq data analysis by attenuating dataset-specific noise and batch effects, enabling more accurate biological inference. Operating at the sample level, this approach overcomes limitations of existing batch correction methods reliant on replicates, demonstrating marked resilience and wider applicability across diverse data types. By facilitating meaningful insight even when replicates are unavailable, the denoising step addresses significant challenges pervasive in bulk RNA-seq inquiry (Fig5).

## Data availability

The scRNA-seq datasets used in the current study can be found in NCBI’s Gene Expression Omnibus and are accessible through the following GEO accession number: GSE60361, GSE85066, GSE109774, GSE118767, GSE84133, GSE158493 and GSE81608; in EBI ArrayExpress are accessible through the following E-MATB accession number: E-MTAB-2805 and E-MTAB-5061; in SRA(Sequence ReadArchive) and is accessible through SRP073767. Bladder bulk RNA-seq data is provided in “bladderbatch” R package (40). TCGA HNSC bulk RNA-seq data can be downloaded directly via https://xenabrowser.net/datapages/ (25)

## Code availability

Source codes implemented can be found at https://github.com/Lan-lab/COMSE

## Acknowledgments

We thank the useful comments on model implementation very much from Yu Yongzhen, MD/Ph.D. candidate, Tsinghua University. This work was supported by Alibaba Group through Alibaba Innovative Research (AIR) Program.

## References

1. Tang, F., Barbacioru, C., Wang, Y., Nordman, E., Lee, C., Xu, N., Wang, X., Bodeau, J., Tuch, B.B., Siddiqui, A., et al. (2009) mRNA-Seq whole-transcriptome analysis of a single cell. Nat Methods, 6, 377–382.

2. Wang, Z., Gerstein, M. and Snyder, M. (2009) RNA-Seq: A revolutionary tool for transcriptomics. Nat Rev Genet, 10, 57–63.

3. Wapinski, O.L., Vierbuchen, T., Qu, K., Lee, Q.Y., Chanda, S., Fuentes, D.R., Giresi, P.G., Ng, Y.H., Marro, S., Neff, N.F., et al. (2013) Hierarchical mechanisms for direct reprogramming of fibroblasts to neurons. Cell, 155, 621.

4. Lee, Y.G., Guruprasad, P., Ghilardi, G., Pajarillo, R., Sauter, C.Tor., Patel, R., Ballard, H.J., Hong, S.Jae., Chun, I., Yang, N., et al. (2022) Modulation of BCL-2 in both T Cells and Tumor Cells to Enhance Chimeric Antigen Receptor T cell Immunotherapy against Cancer. Cancer Discov, 10.1158/2159-8290.cd-21-1026.

5. Otero-Garcia, M., Mahajani, S.U., Wakhloo, D., Tang, W., Xue, Y.Q., Morabito, S., Pan, J., Oberhauser, J., Madira, A.E., Shakouri, T., et al. (2022) Molecular signatures underlying neurofibrillary tangle susceptibility in Alzheimer’s disease. Neuron, 10.1016/j.neuron.2022.06.021.

6. Shalek, A.K., Satija, R., Shuga, J., Trombetta, J.J., Gennert, D., Lu, D., Chen, P., Gertner, R.S., Gaublomme, J.T., Yosef, N., et al. (2014) Single-cell RNA-seq reveals dynamic paracrine control of cellular variation. Nature, 510, 363–369.

7. Wu, A.R., Neff, N.F., Kalisky, T., Dalerba, P., Treutlein, B., Rothenberg, M.E., Mburu, F.M., Mantalas, G.L., Sim, S., Clarke, M.F., et al. (2014) Quantitative assessment of single-cell RNA-sequencing methods. Nat Methods, 11, 41–46.

8. Hashimshony, T., Wagner, F., Sher, N. and Yanai, I. (2012) CEL-Seq: Single-Cell RNA-Seq by Multiplexed Linear Amplification. Cell Rep, 2, 666–673.

9. Islam, S., Kjällquist, U., Moliner, A., Zajac, P., Fan, J.B., Lönnerberg, P. and Linnarsson, S. (2011) Characterization of the single-cell transcriptional landscape by highly multiplex RNA-seq. Genome Res, 21, 1160–1167.

10. Andrews, T.S. and Hemberg, M. (2019) M3Drop: Dropout-based feature selection for scRNASeq. Bioinformatics, 35, 2865–2867.

11. Macosko, E.Z., Basu, A., Satija, R., Nemesh, J., Shekhar, K., Goldman, M., Tirosh, I., Bialas, A.R., Kamitaki, N., Martersteck, E.M., et al. (2015) Highly parallel genome-wide expression profiling of individual cells using nanoliter droplets. Cell, 161, 1202–1214.

12. Hicks, S.C., Townes, F.W., Teng, M. and Irizarry, R.A. (2018) Missing data and technical variability in single-cell RNA-sequencing experiments. Biostatistics, 19, 562–578.

13. Satija, R., Farrell, J.A., Gennert, D., Schier, A.F. and Regev, A. (2015) Spatial reconstruction of single-cell gene expression data. Nat Biotechnol, 33, 495–502.

14. Lun, A.T.L., McCarthy, D.J. and Marioni, J.C. (2016) A step-by-step workflow for low-level analysis of single-cell RNA-seq data [version 1; referees: 5 approved with reservations]. F1000Res, 5.

15. Vallejos, C.A., Marioni, J.C. and Richardson, S. (2015) BASiCS: Bayesian Analysis of Single-Cell Sequencing Data. PLoS Comput Biol, 11.

16. Buettner, F., Natarajan, K.N., Casale, F.P., Proserpio, V., Scialdone, A., Theis, F.J., Teichmann, S.A., Marioni, J.C. and Stegle, O. (2015) Computational analysis of cell-to-cell heterogeneity in single-cell RNA-sequencing data reveals hidden subpopulations of cells. Nat Biotechnol, 33, 155–160.

17. Qiu, X., Mao, Q., Tang, Y., Wang, L., Chawla, R., Pliner, H.A. and Trapnell, C. (2017) Reversed graph embedding resolves complex single-cell trajectories. Nat Methods, 14, 979–982.

18. Blondel, V.D., Guillaume, J.-L., Lambiotte, R. and Lefebvre, E. (2008) Fast unfolding of communities in large networks. Journal of Statistical Mechanics: Theory and Experiment, 10.1088/1742-5468/2008/10/P10008.

19. He, X., Cai, D. and Niyogi, P. (2005) Laplacian Score for Feature Selection. In 18th International Conference on Neural Information Processing Systems.pp. 507–514.

20. DeTomaso, D. and Yosef, N. (2021) Hotspot identifies informative gene modules across modalities of single-cell genomics. Cell Syst, 12, 446–456.e9.

21. Becht, E., McInnes, L., Healy, J., Dutertre, C.A., Kwok, I.W.H., Ng, L.G., Ginhoux, F. and Newell, E.W. (2019) Dimensionality reduction for visualizing single-cell data using UMAP. Nat Biotechnol, 37, 38–47.

22. Sara Aibar and Stein Aerts AUCell: AUCell: Analysis of ‘Gene Set’ Activity in Single-Cell RNA-Seq Data.

23. Rousseeuw, P.J. (1987) Silhouettes: A graphical aid to the interpretation and validation of cluster analysis. J Comput Appl Math, 20, 53–65.

24. Dyrskjøt, L., Kruhøffer, M., Thykjaer, T., Marcussen, N., Jensen, J.L., Møller, K. and Ørntoft, T.F. (2004) Gene Expression in the Urinary Bladder: A Common Carcinoma in Situ Gene Expression Signature Exists Disregarding Histopathological Classification.

25. Goldman, M.J., Craft, B., Hastie, M., Repečka, K., McDade, F., Kamath, A., Banerjee, A., Luo, Y., Rogers, D., Brooks, A.N., et al. (2020) Visualizing and interpreting cancer genomics data via the Xena platform. Nat Biotechnol, 38, 675–678.

26. Ritchie, M.E., Phipson, B., Wu, D., Hu, Y., Law, C.W., Shi, W. and Smyth, G.K. (2015) Limma powers differential expression analyses for RNA-sequencing and microarray studies. Nucleic Acids Res, 43, e47.

27. McDavid, A., Finak, G. and Gottardo, R. (2016) The contribution of cell cycle to heterogeneity in single-cell RNA-seq data. Nat Biotechnol, 34, 591–593.

28. Rapsomaniki, M.A., Lun, X.-K., Woerner, S., Laumanns, M., Bodenmiller, B. and Martínez, M.R. (2018) CellCycleTRACER accounts for cell cycle and volume in mass cytometry data. Nat Commun, 9, 632.

29. Zeisel, A., Muñoz-Manchado, A.B., Codeluppi, S., Lönnerberg, P., la Manno, G., Juréus, A., Marques, S., Munguba, H., He, L., Betsholtz, C., et al. (2015) Cell types in the mouse cortex and hippocampus revealed by single-cell RNA-seq. Science (1979), 347, 1138–1142.

30. Baron, M., Veres, A., Wolock, S.L., Faust, A.L., Gaujoux, R., Vetere, A., Ryu, J.H., Wagner, B.K., Shen-Orr, S.S., Klein, A.M., et al. (2016) A Single-Cell Transcriptomic Map of the Human and Mouse Pancreas Reveals Inter- and Intra-cell Population Structure. Cell Syst, 3, 346–360.e4.

31. Bunis, D.G., Bronevetsky, Y., Krow-Lucal, E., Bhakta, N.R., Kim, C.C., Nerella, S., Jones, N., Mendoza, V.F., Bryson, Y.J., Gern, J.E., et al. (2021) Single-Cell Mapping of Progressive Fetal-to-Adult Transition in Human Naive T Cells. Cell Rep, 34.

32. Duò, A., Robinson, M.D. and Soneson, C. (2020) A systematic performance evaluation of clustering methods for single-cell RNA-seq data. F1000Res, 7, 1141.

33. Koh, P.W., Sinha, R., Barkal, A.A., Morganti, R.M., Chen, A., Weissman, I.L., Ang, L.T., Kundaje, A. and Loh, K.M. (2016) An atlas of transcriptional, chromatin accessibility, and surface marker changes in human mesoderm development. Sci Data, 3, 160109.

34. Kuhn, S., Gritti, L., Crooks, D. and Dombrowski, Y. (2019) Oligodendrocytes in development, myelin generation and beyond. Cells, 8.

35. Jovic, D., Liang, X., Zeng, H., Lin, L., Xu, F. and Luo, Y. (2022) Single-cell RNA sequencing technologies and applications: A brief overview. Clin Transl Med, 12.

36. Hicks, S.C., Townes, F.W., Teng, M. and Irizarry, R.A. (2018) Missing data and technical variability in single-cell RNA-sequencing experiments. Biostatistics, 19, 562–578.

37. Xin, Y., Kim, J., Okamoto, H., Ni, M., Wei, Y., Adler, C., Murphy, A.J., Yancopoulos, G.D., Lin, C. and Gromada, J. (2016) RNA Sequencing of Single Human Islet Cells Reveals Type 2 Diabetes Genes. Cell Metab, 24, 608–615.

38. Segerstolpe,Å., Palasantza, A., Eliasson, P., Andersson, E.-M., Andréasson, A.-C., Sun, X., Picelli, S., Sabirsh, A., Clausen, M., Bjursell, M.K., et al. (2016) Single-Cell Transcriptome Profiling of Human Pancreatic Islets in Health and Type 2 Diabetes. Cell Metab, 24, 593–607.

39. Johnson, W.E., Li, C. and Rabinovic, A. (2007) Adjusting batch effects in microarray expression data using empirical Bayes methods. Biostatistics, 8, 118–127.

40. Jeffrey T. Leek bladderbatch: Bladder gene expression data illustrating batch effects.

