## Supplementary Information for "COMSE: Analysis of Single-Cell RNA-seq Data Using Community Detection Based Feature Selection"

**Simulation experiments in FigS1**

The simulated data comprised three groups with three subsets of features; each group contained 100 samples. Subset 1 contained 100 features drawn from three normal distributions with small means and variances corresponding to the different groups, with the mean differences between the three normal distributions also small. Subset 2 contained 100 features drawn from three normal distributions corresponding to the different groups, but with larger means and variances and larger differences between the means of the three distributions. Subset 3 contained 300 features drawn from a single normal distribution irrespective of group, with the largest variance, as follows.

For features in subset1:

$$\mu_{ij} \sim U(2{,}2.1), g_{ij} \sim N(\mu_{ij}, 0.25)$$

where i, j are indexes for group and gene.

For features in subset2:

$$\mu_{ij} \sim U(8{,}10), g_{ij} \sim N(\mu_{ij}, 1)$$

where i, j are indexes for group and gene.

For features in subset3:

$$\mu_j \sim U(8{,}10), g_j \sim N(\mu_{ij}, 2)$$

where j is index for gene.

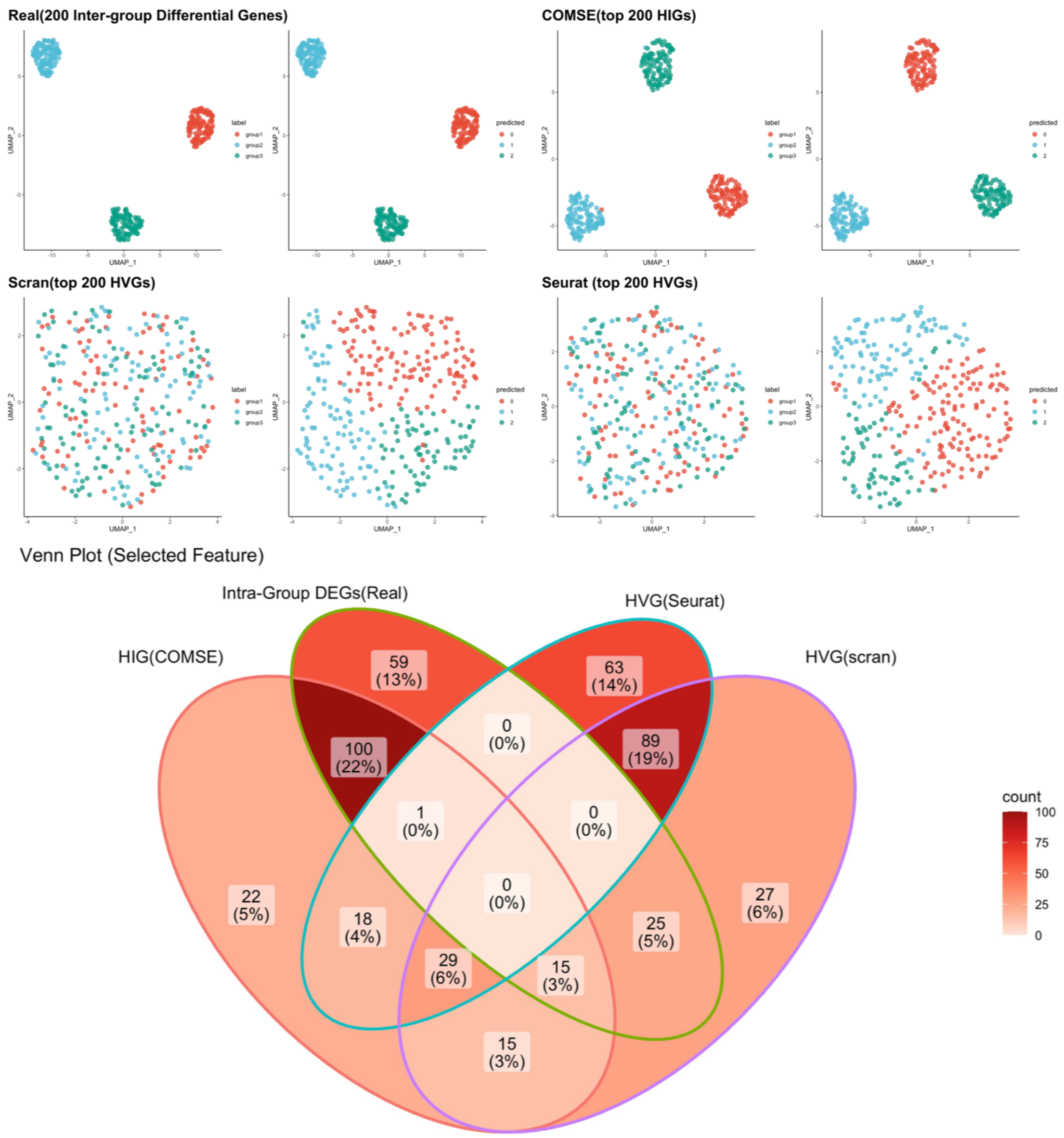


**FigS1.** The UMAP plots of the simulated data contained three group with 500 features, each group contained 100 samples labelled by predicted label using 200 inter-group differentially expressed genes and top 200 genes selected by different feature selection methods and true label. The Venn plots of inter-group differentially expressed genes and top 200 genes selected by different feature selection methods

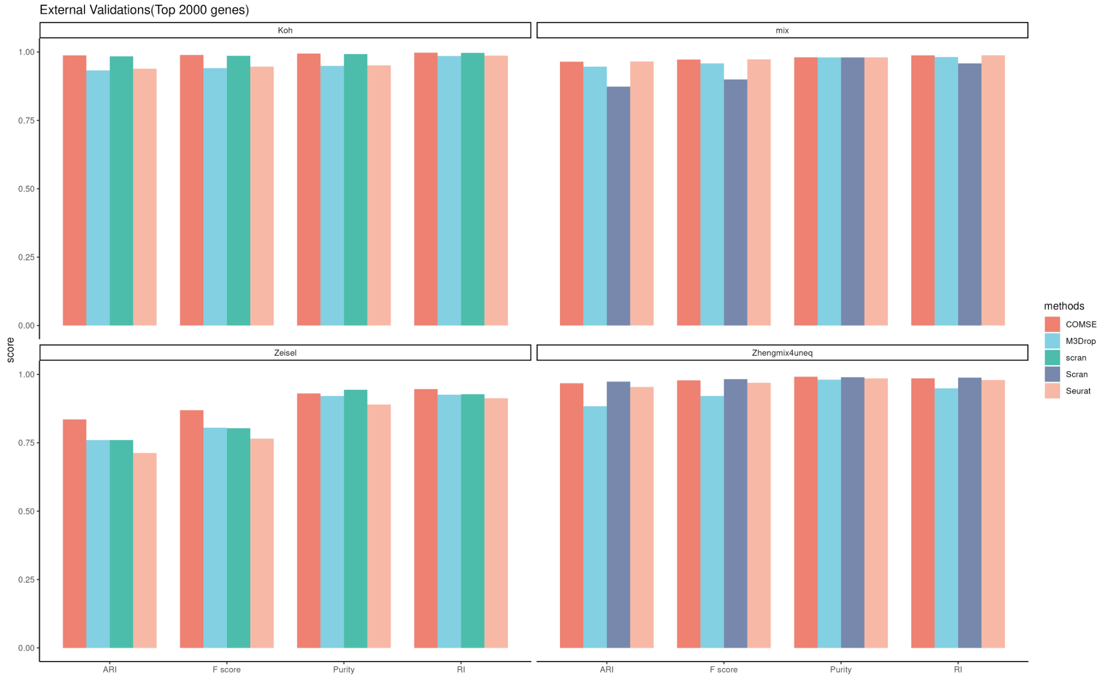


**Figure S2.** COMSE yields more accurate cell classification results with four scRNA-seq datasets. The cell clustering results using top 2,000 genes selected by COMSE only and other three widely used HVG selection methods of three heterogeneous public scRNA-seq datasets with true labels, including an unequal mixture of B cells, naive cytotoxic T-cells, CD14 monocytes and regulatory T-cells (Zhengmix4uneq) dataset, mouse brain cortex (Zeisel) dataset, the mixture of five human lung adenocarcinoma cell lines (mixology10x5cl) dataset and purified H7 human embryonic stem cells at different developmental stages (Koh) dataset. Four external clustering validations (F score, CSI, PSI, ARI, and NMI) were used to indicate the performance and robustness of the five methods on four public scRNA-seq datasets.

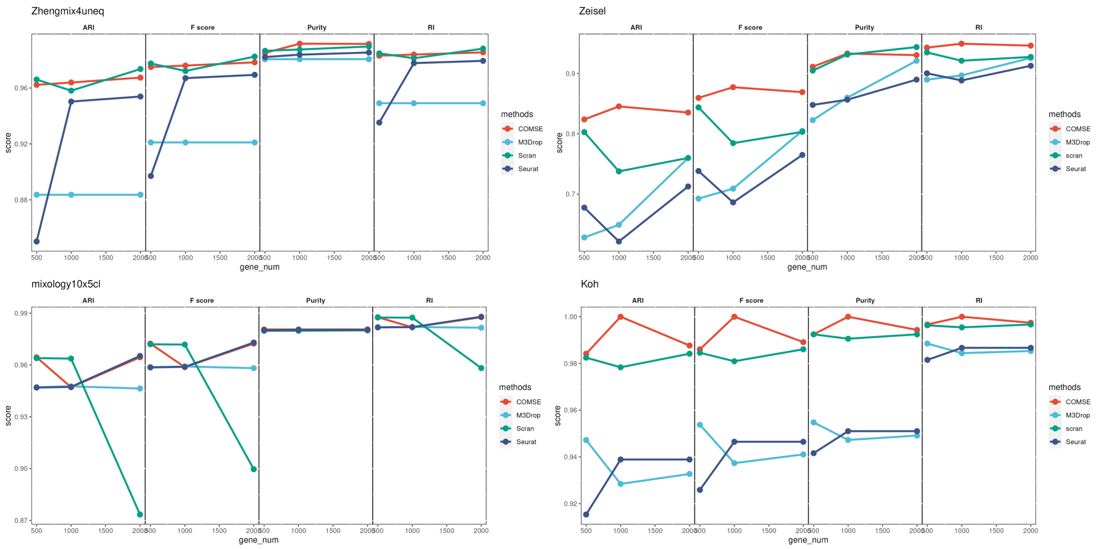


**Figure S3.** COMSE yields more stable cell classification results with four scRNA-seq datasets. The cell clustering results using top 500, 1,000, and 2,000 genes selected by COMSE and other four widely used HVG selection methods of four public scRNA-seq datasets with solid true labels, including an unequal mixture of B cells, naive cytotoxic T-cells, CD14 monocytes and regulatory T-cells (Zhengmix4uneq) dataset, mouse brain cortex (Zeisel) dataset, the mixture of five human lung adenocarcinoma cell lines (mixology10x5cl) dataset and purified H7 human embryonic stem cells at different developmental stages (Koh) dataset. Four external clustering validations (F score, CSI, PSI, ARI, and NMI) were used to indicate the performance and robustness of the five methods on four public scRNA-seq datasets

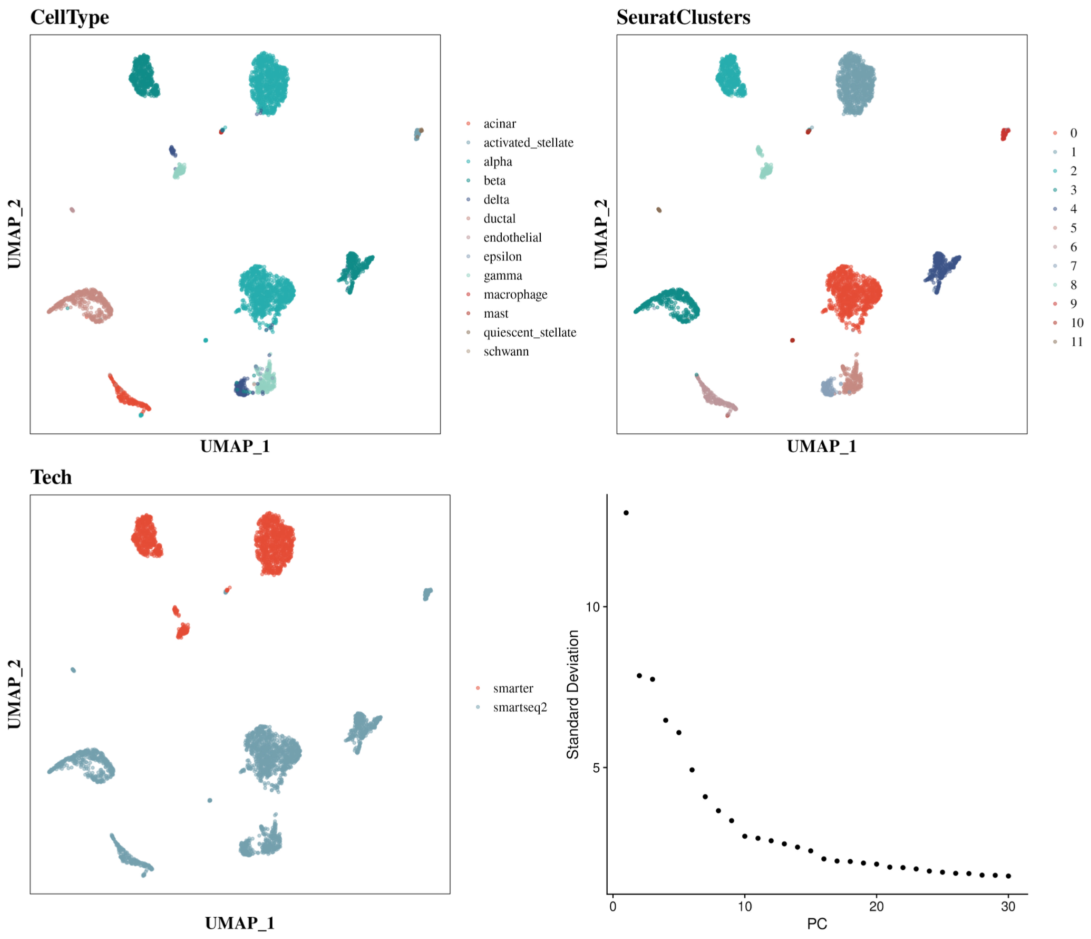


**Figure S4.** UMAP plots of the human pancreatic islet datasets labelled by cell types, predicted label using Seurat, scRNA-seq protocols. We extracted 2000 highly variable genes for downstream clustering. The first 10 PCA components of the data matrix (Seurat) as the input for UMAP dimension reduction and 2D visualization. The best n PCA components used in Seurat were selected based on the Elbow plot.

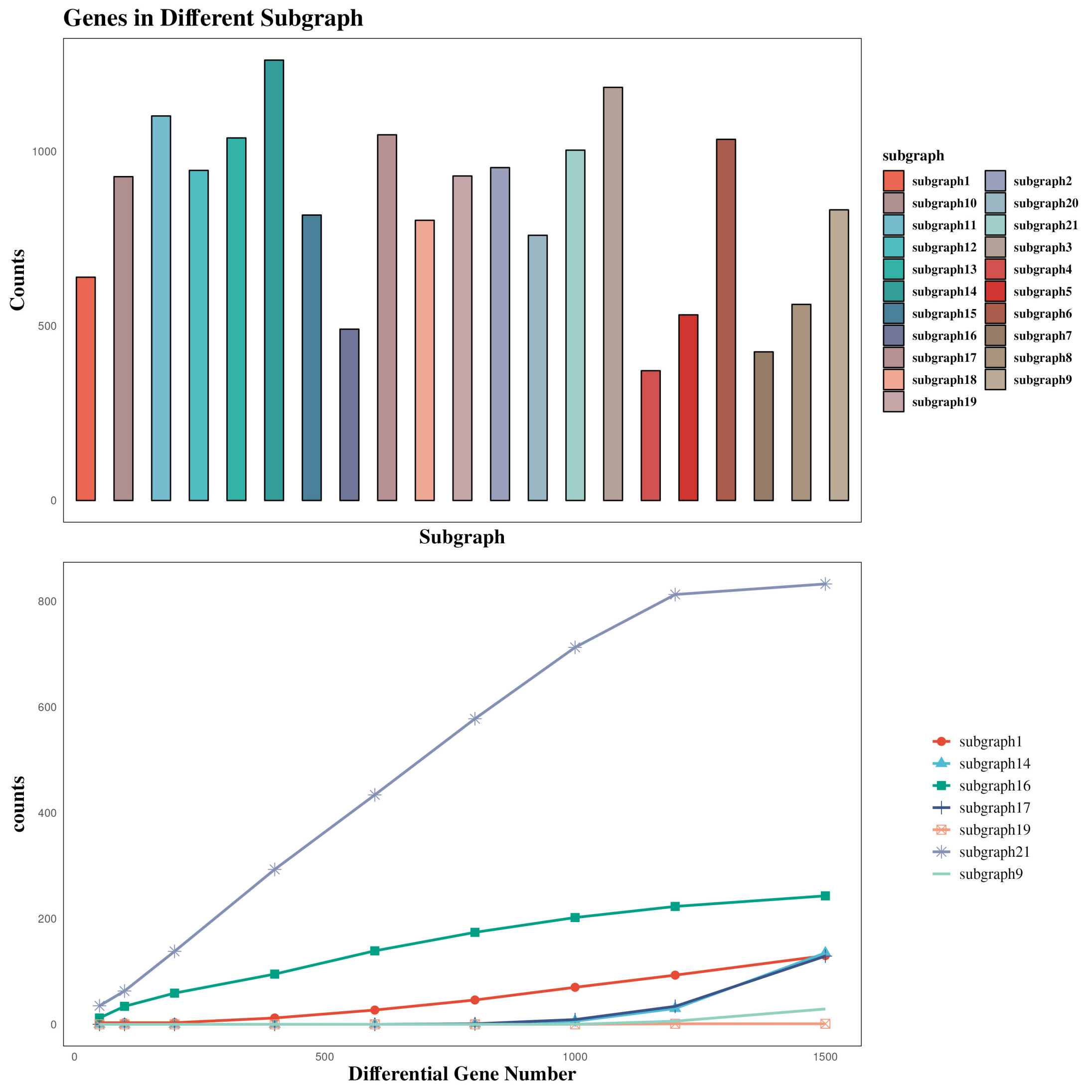


**Figure S5.** The top bar graph shown the absolute gene number in each subgraph estimated by community detection algorithm in COMSE, and the bottom line graph shows the number of top n differentially expressed gene between two scRNA-seq protocols (smater versus smartseq2) using FindMarkers provided by Seurat in each subgraph.

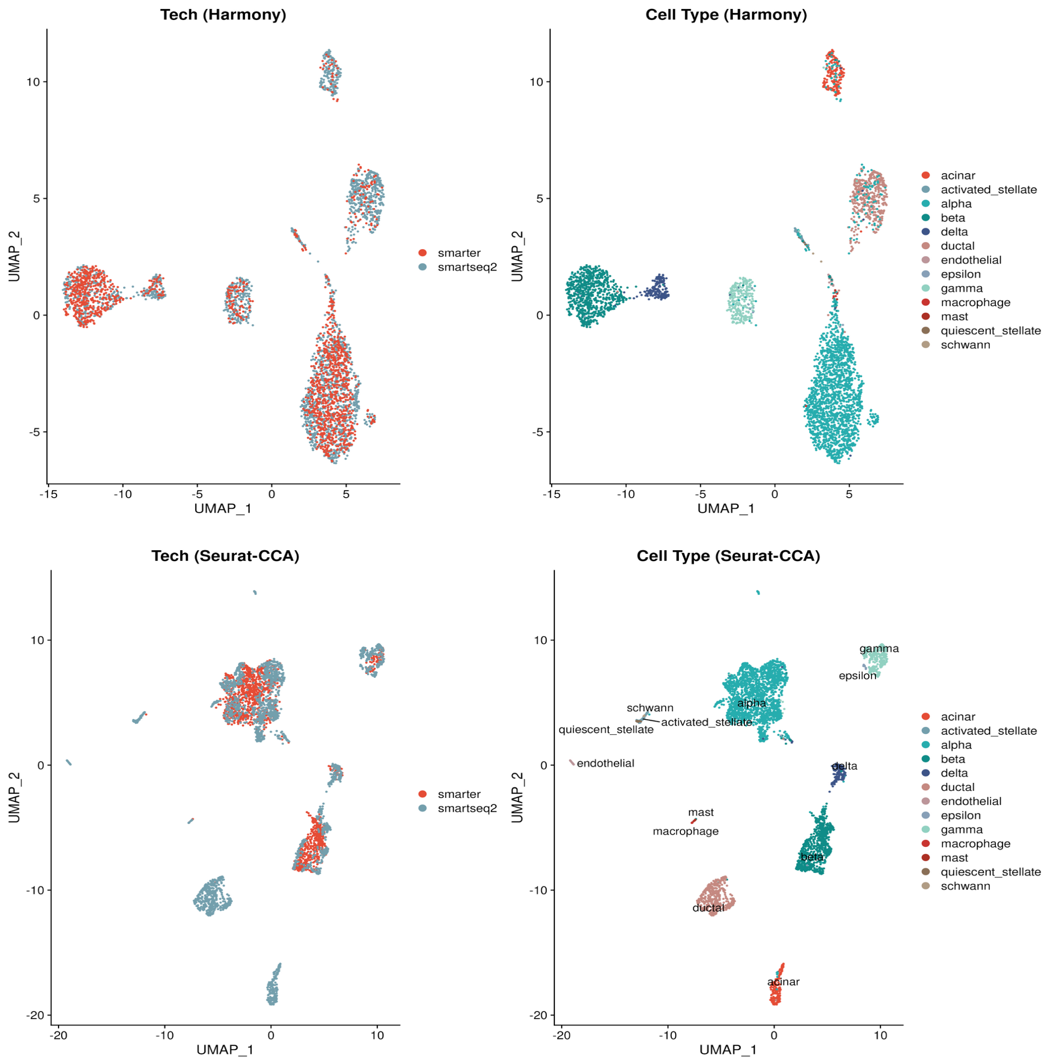


**Figure S6.** The result of commonly used batch effect removal methods. After using the Harmony and CCA provided by Seurat respectively, UMAP plots of the human pancreatic islet datasets obtained with top 2000 genes selected by COMSE labelled by scRNA-seq protocols and cell types.

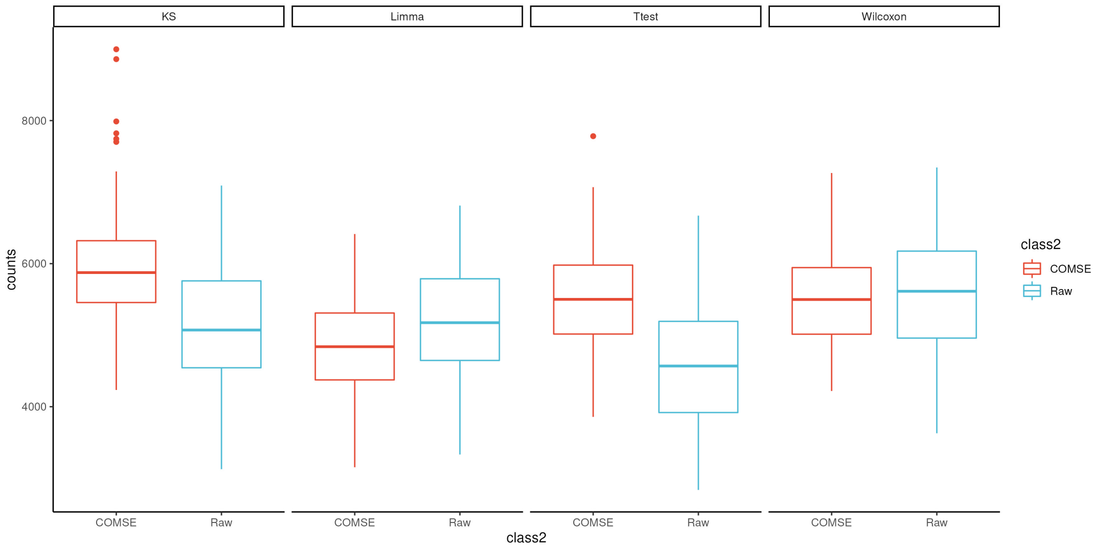


FigS7. Boxplots of the counts of DEGs screened via each approach including t-test, Wilcoxon-test, limma and KS-test using original and denoised down-sampled data collections.
